# Development of nascent focal adhesions in spreading cells

**DOI:** 10.1101/2020.03.03.972992

**Authors:** Neil Ibata, Eugene M. Terentjev

## Abstract

Cell spreading provides one of the simplest configurations in which eukaryotic cells develop angular symmetry-breaking assemblies of mechanosensing and mechanotransducive organelles in preparation for cell differentiation and movement. By identifying the edge of the cell-ECM adhesion area as having an important role in mechanosensor complex aggregation, we consider the spatial patterns arising on this edge, within a 1D lattice model of the nearest-neighbour interaction between individual integrin-mediated mechanosensors. We obtain the Ginzburg-Landau free energy for this model and analyse the spectrum of spatial modes as the cell spreads and increases the contact area. We test the plausibility of our model by comparing its predictions for the azimuthal angular frequency of aggregation of mechanosensors into nascent focal adhesions (FAs) to observations of the paxillin distribution in spreading fibroblasts.

**STATEMENT OF SIGNIFICANCE:** The topic of cell adhesion on substrates is very active, with numerous theoretical, experimental and computer simulation studies probing the mechanisms and signalling pathways of cell response to interacting with substrate. Integrin-based adhesion complexes are known to be the individual units of this process, and their dense aggregation into focal adhesions leads to cells developing asymmetry, polarity, and eventually - locomotion. Here we develop a theoretical model that suggests that physical interactions between individual adhesion complexes is the factor that defines the initial breaking of symmetry of the cell spreading on substrate, and predicts the characteristic wavelength of modulation above the critical size of adhesion area.

## 1 Introduction

Most eukaryotic cells respond to external physical stimuli, and exert forces on surrounding cells and the extracellular matrix (ECM) by substantially changing their global shape and developing localised mechanosentive and mechan-otransducive organelles – notably, focal adhesions (FAs), stress fibres and motile membrane protrusions (1, 2). These morphological changes can be rapid and may help to determine cell function, including differentiation (3), motility (4, 5) and proliferation (6).

When a cell is located in an inhomogeneous medium with external physical and chemical cues, its shape becomes less symmetric in response (7). Well-known examples include chemotaxis (8) and durotaxis (9, 10), i.e. the cell’s response to gradients of chemical concentration and the ECM stiffness, respectively, in which the eukaryotic cell takes on a characteristic polarized shape in preparation for directed motion in the direction of the gradient.

Although molecular pathways for both of these mechanisms are currently under investigation (11, 12), and a full network of interactions between the known components of the integrin adhesome underpinning durotaxis has been elucidated (13), a detailed understanding of how their individual components arrange themselves spatially in order to lay down the necessary asymmetries in the cell is still lacking. It is clear that a valid physical model accounting for the assembly and function of the force sensing and force transduction machinery of a generic cell is required in order to progress with this problem.

Attempting such an analysis in a general geometry with a complex set of external cues is very complicated, and one must begin by reducing the problem down to the simplest possible (while experimentally viable) configuration which still displays a defined organisation of the constituent molecules within these pathways.

The mechanosensitive and mechanotransducive machinery has been found to assemble the same organelles both in the presence and absence of large scale physical gradients (*e.g.* of stiffness and force); moreover, cells placed in a homogeneous medium can still undergo stiffness-induced differentiation and, more remarkably, they can change their shape in such a manner as to allow for short periods of straight-line motion as demonstrated e.g. by Hartman et al. (14). The clearest example of this comes from one of the simplest possible examples of cell-matrix interaction. When a near-spherically symmetric cell is taken from suspension and placed on an isotropic and homogeneous 2D substrate, its radial shape profile changes drastically in a process known as cell spreading due to a combination of energetically favourable surface adhesion and costly remodelling of its cytoskeleton (15). The change in the overall shape of the cell has been modelled using hydrodynamic arguments: both the initial fast, passive spreading phase (first Bruinsma and Sackmann (16), and then Cuvelier et al. (17) model the cell as a viscoelastic vesicle) and the subsequent slower, active (energetically costly) spreading phase (18–20).

In conjunction with the flattening of the height profile, an azimuthal symmetry-breaking pattern of local cytoskeleton-driven membrane protrusions develops in the plane of ECM attachment, including lamellipodia, filopodia (21) and fibronexi (22). In polarised cells, these pseudopodia are necessary for motility, and define the mode of cell motion (only a subset of the full possible modes of motion described in 3D media by Petrie and Yamada (23) actually applies to the motion of a cell placed on a 2D ECM substrate). But these protrusions themselves were found to mostly form towards the end of cell spreading (Johnston et al. (24) observed that lamellipodia only developed in fibroblasts after 30 minutes of cell spreading on fibronectin), that is to say after the time at which cell-matrix integrin-mediated adhesion receptors which mediate ECM-cytoskeleton interactions aggregate into an obviously heterogeneous distribution. Key experiments by Fouchard et al. (25), and others (26) suggest that this takes place ataround 3 minutes after initial contact in a similar set-up.

A detailed explanation for the formation of membrane protrusions is complex, and possible mechanisms have been examined in some detail, in particular in moving cells (28, 29). A theoretical model for this appears currently out of reach, but in light of current models of filopodia formation in terms of nascent focal adhesions (30, 31), we suggest that a first step towards explaining the large-scale organisation of the outer cytoskeleton of the cell on substrate involves accounting for the evolution of aggregates of indivudual membrane-bound force sensors into the observed distribution of nascent adhesions.

In this endeavour, time-lapse immunofluorescence imaging of the paxillin distribution within spreading fibroblasts on fibronectin-coated glass (25, 26) have proven invaluable. Their experiments highlight some key elements of this initial angular symmetry-breaking sensor concentration. In particular: integrin-containing nascent focal adehsions (FAs) first develop only once the cell has reached a particular shape, initially develop very close to the edge of the cell-ECM contact area, and appear to aggregate on a specific length scale around the edge. The scenario is shown schematically in Fig. 1.

**Figure 1:**
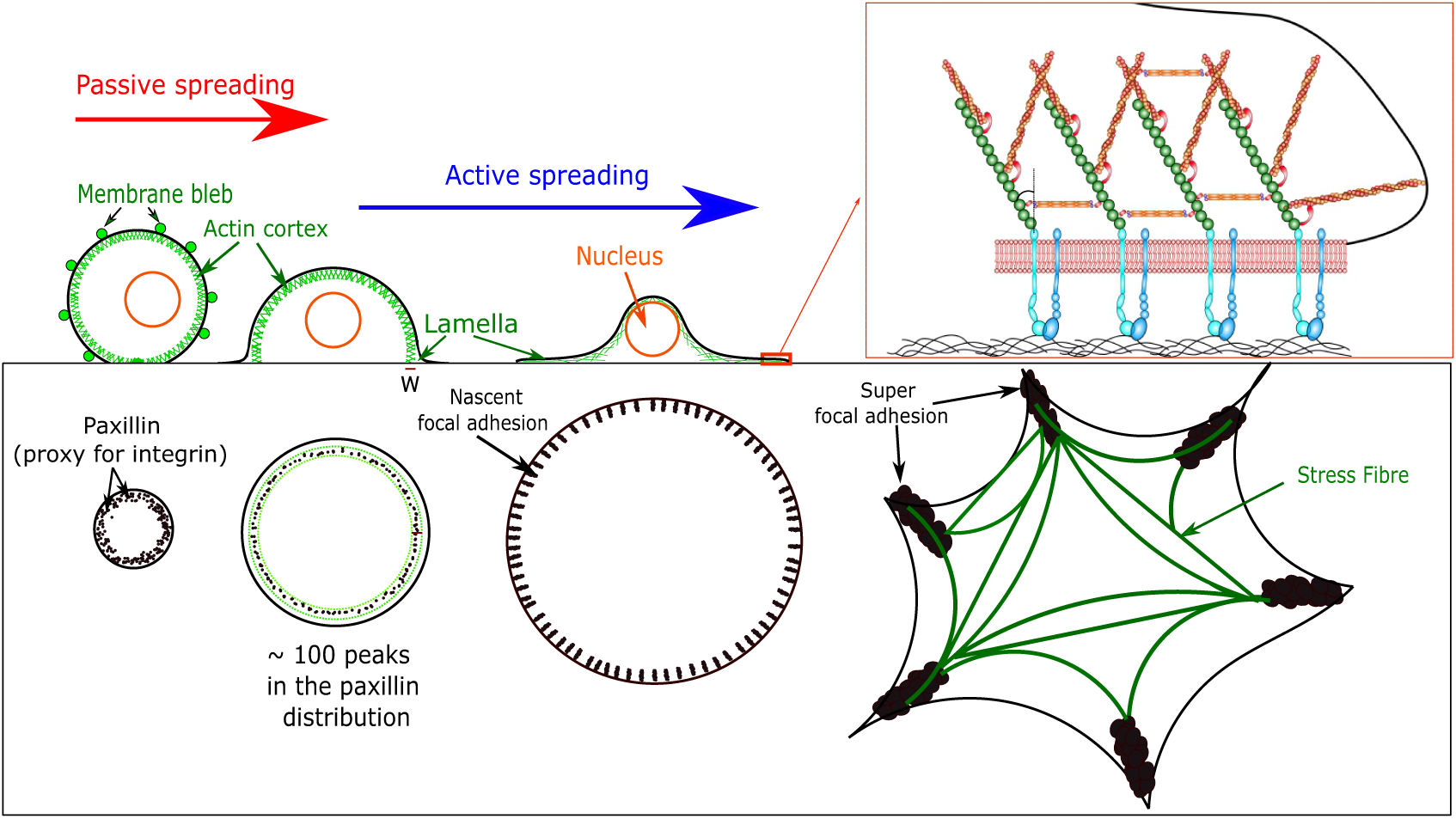
Upper panels: side-on sketch of a cell spreading on a 2D ECM surface. The cell settles by sedimentation, passive spreading and then active spreading (27). This results in a balance between adhesion energy and acto-myosin cortex deformation at first, followed by a protrusive force exerted by F-actin polymerisation and transduced at focal adhesions (inset on the right). Lower panels: a view from above on the increasing cell-ECM contact area during spreading. As observed by Fouchard et al. (25), mechanosensors aggregate into FAs during cell spreading. After cell spreading, on sufficiently stiff substrates, FAs may aggregate further into super-FAs at fibronexi connected by large actin stress fibres (rightmost sketch).

The development of spatial patterns of aggregates of cell-surface receptors has been analytically examined in related paradigms by Guo and Levine (32, 33). In that work, simple nearest-neighbour interactions between sensors arranged as a lattice gas were shown to lead to a phase transition at which a uniform sensor distribution starts to aggregate into macroscopic clumps at a certain temperature, sensor concentration and sensor-sensor interaction energy.

In the case of cell spreading, a lattice-model based analysis of this kind is complicated by the expansion of the cell-ECM contact area, which continuously increases the size of the lattice, and not only changes the sensor concentration but also the number of possible arrangements of the sensor distribution. However, a phase transition within the integrin-mediated sensor distribution modelled as a lattice gas is also possible in a configuration in which the cell spreads. Here, both the size of the contact area and the interaction energy change during the spreading process and could both potentially act as a ‘temperature’-like parameter in an Ising-like analysis of the phase transition between the uniform and clustered sensor distributions.

The spreading process not only changes the conditions for the structural phase transition to occur, but also changes the subsequent evolution of the angular pattern of sensors. Whereas a specific fastest-growing spatial mode may clearly be identified for a phase transition for which the critical parameter does not change markedly after the transition, the continuing cell spreading complicates the situation at hand. As we will see, one needs to carefully examine the time at which specific spatial modes begin to grow, as well as their growth time, in order to at least qualitatively examine the spatial spectrum of angular aggregates of integrin-mediated mechanosensors. Finally, this will need to be compared with results from immunofluorescence imagery of spreading cells.

### 1.1 A physical approach is known to help explain large-scale cell shape changes during cell spreading

The shape of a eukaryotic cell in planktonic suspension is highly constrained to small thermal fluctuations about a spherically symmetric shape if its membrane is placed under sufficient tension, by the combined effect of the acto-myosin cortex and bleb formation (15). Assuming that a tense giant unilamellar vesicle (GUV) is a good model for a cell in suspension, measurements (34) suggest that typical membrane fluctuations are less than 25*nm* in size. In contrast, once the cell adheres to a surface, Monzel and Sengupta (35) report that thermal membrane fluctuations are even smaller (dying down to the sub-*nm* scale) and can be ignored.

At a similar length scale, a large number of transient actin spikes protrude out from the cell body, as seen at the very start of cell spreading in electron microscope observations of fibroblasts by Albuschies and Vogel (36). However, given their very small size and because they are clearly under no load and therefore cannot transmit any force into the cell’s intracellular force-sensing machinery, they most likely do not substantially contribute to the cell’s response to physical stimuli.

When the cell comes in contact with a plane surface covered with ECM proteins, a small portion of the cell initially anchors to the substrate, possibly by hydrophobic or Van der Waals interactions (16, 37); this corresponds to Phase I of spreading as classified by Khalili and Ahmad (27). The cell membrane contains a large number of receptors which participate in cell-ECM ligand binding (38). These then provide the cell with a surface tension, which is an energy source for the deformation of the acto-myosin cortex that takes place at the start of cell spreading. This process does not require the cell to expend any energy in the form of ATP-dependent protein polymerisation or motor action of myosin cross-linking, and we will thus refer to this initial period as ‘passive spreading’, which corresponds to Phase II of cell spreading following the aforementioned classification (27).

On sufficiently stiff substrates, this is subsequently followed by an ‘active spreading’ phase (20), in which the energetically costly polymerisation of new F-actin provides the necessary outwards force to break cross-linkers or F-actin filaments within the acto-myosin cortex. This phase lasts longer and involves slower spreading until the cell reaches its maximal area, and corresponds to Phase III of spreading (27). The duration of these processes depends heavily on temperature, i.e. the internal thermally-activated processes, as shown by Bell et al. (39).

These phases have been shown to conform to simple physical hydrodynamic arguments: passive spreading matches well to the spreading of a composite viscous drop as shown by Cuvelier et al. (17) and Étienne and Duperray (40), whereas the active spreading can be understood as a combination of viscous dissipation due to the partial disassembly of the acto-myosin cortex and the flow of cell material due to F-actin polymerisation (18).

A physical reasoning is very useful in analysing the evolution of the overall shape of a cell in contact with a uniform flat ECM surface. But an analytical approach is also well suited to examine the development of longrange supramolecular order necessary for the development of aggregates of mechanosensitive molecules from short range interactions due to the high degree of symmetry which characterises ideal cell spreading.

There is a caveat to these symmetry considerations which somewhat complicates an analytical exploration. As evidenced by atomic force microscopy measurements of spreading cells, from suspension to full attachment, a eukaryotic cell maintains its membrane tension and stores excess membrane materials in suspension in a series of invaginations and membrane blebs (15). These are slowly released during cell spreading, and result in fluctuations in acto-myosin cortex tension and help explain transient protrusions of the contact area. These protrusions are much larger than the lengthscale for sensor aggregation (with typical bleb radii *r*_bleb_ > 1*µm*), suggesting that their local distortion on the cell shape is sufficiently small as to not substantially affect the geometry in which the aggregation of sensors takes place.

Because thermal fluctuations have a negligible effect on the shape of the cell during spreading, it seems likely that rapid bleb retraction is the main mechanism which generates a noisy cell-ECM contact area. This takes place on the timescale of less than a minute as observed by Charras et al. (41). In addition, the signature of bleb-induced noise is very different to thermal noise, leading to large *µm*-scale membrane fluctuations (40).

### 1.2 Interactions between assembled integrin-mediated adhesions are key to forming azimuthal patterns

The typical radial and vertical shapes of spreading cells have been accurately modelled using a hydrodynamic approach. But the development of the in-plane pattern of membrane protrusions and mechanosensitive organelles observed in many fully-spread cells has not yet been properly examined. This angular shape is often essential for biological function: for instance, on high stress matrices, fibroblasts may differentiate into myofibroblasts in order to participate in fibrosis and wound healing (42, 43). These cells display a small number of super-focal adhesions (44), large dense clusters of integrin-mediated adhesions which help the cell set up a set of long-lived acto-myosin stress fibres and maintain a large tension throughout and a typical ‘star-like’ shape (45), as illustrated in Fig. 1.

An initial angular shape pattern is laid down prior to these large-scale developments, during the spreading process itself. Cell-matrix adhesions, as noted above, play an important role in the development of a global combined radial/vertical cell shape, likely also play a key role in the development of such an angular pattern.

Although there are many different types of adhesions in cells, one of the most prevalent and best studied are the integrin-mediated complexes (46, 47) involved in coupling the actin cytoskeleton to the ECM substrate. This coupling sets these protein complexes and their aggregates in FAs apart from other types of adhesions, as it allows for dynamic actin filaments to push (and pull) against the adhesion. This will result in a local force passed across the cell membrane to the substrate, and in turn in an observed overall angular shape pattern in the cell as seen from above the adhesion plane.

Macroscopic focal adhesions seen in the fully spread cell are large aggregates of transmembrane integrin complexes bound to a large number (over 50) of cytoplasmic proteins which have been largely identified in immunofluorescence staining experiments of cells with well-developed FAs (48). Notably, vinculin, talin, paxillin, zyxin, *α*-actinin, VASP, FAK, phosphotyrosine proteins, and integrin have all been shown to be found in large quantities in nascent and fully-developed focal adhesions (49).

The structures and binding possibilities of these component proteins have been thoroughly investigated. This wealth of data strongly suggests that each individual integrin interacts with a cytoplasmic complex, which serves both as a force transduction chain and as a source for force-induced signalling (50, 51). In this picture, the membrane-bound integrin and the associated cytoplasmic plaque form an individual mechanosensing complex. If most if not all of the observed proteins associated with the cytoplasmic domain of the mechanosensing complex are present in each mechanosensor, the lateral size of the mechanosensor – and therefore the distance between neighbouring integrins in a developed densely-packed FA – is limited by the size of its constituent proteins.

When the mechanosensor complexes are fully developed, the distribution of each of the constituent proteins found within the cytoplasmic plaque may be used as a proxy for the actual mechanosensor and integrin distribution. But in the case of cell spreading, this reasoning can be taken a little further: when the cell is in planktonic suspension, the constituent proteins of mechanosensors have not been found to have any spatial structure. Indeed Mueller et al. (52) found that the concentration of the key protein talin is diffuse within the cell in suspension, while the concentration of integrin is uniformly spread across the cell membrane. This implies that at the start of cell spreading, the concentration of the components of the cytoplasmic domain of the mechanosensor near the cell membrane will be quite small, and so one expects to be able to use such proxy measurements to assess the development and growth of the number of full mechanosensing complexes – by examining the intensity of paxillin, talin etc. in the immediate vicinity of the membrane.

Indeed, observations of paxillin distribution in fibroblasts by Fouchard et al. (25) showed that an azimuthal spatial pattern with a specific aggregation length scale developed near the edge of the growing contact area after ≈ 3 minutes of spreading. For the cells that they studied, this corresponded to a configuration in which the acto-myosin cortex developed in suspension had been distorted beyond a hemispheric configuration (results by Cuvelier et al. (17) suggest that this time is substantially after the end of passive spreading). As far as we are aware, this corresponds to the first time at which an angular structure develops in the spreading cell, so it seems likely that attractive interactions between mechanosensor complexes are directly responsible for the observed aggregation.

The component proteins of a mechanosensor complex take a characteristic time to be recruited to the membrane sites, depending on a (fast) diffusion time, and a (much slower) binding time. By examining population dynamics of the time at which the cell becomes hemispherical in shape, Bell et al. (39) found that a 5^th^ order power law determined the number of spread cells at a particular early spreading time, suggesting 5 slow thermally-activated steps in the complex assembly. Once in the hemispherical shape configuration, the spreading cell is already in the active spreading phase in which F-actin polymerisation in the lamella provides the required protrusive force. One explanation for this is that the rate-limiting factor for fast spreading to begin is the formation of the full mechanosensor complex (53), which would then involve four slow kinetic steps (rate dependent on temperature). The fully-assembled mechanosensing complex would then positively signal for Rho/Rac signalling and lead to an activation of F-actin polymerisation. An adaptation of the mechanism of Bell et al. (39) to account for this behaviour is shown in Fig. 2.

**Figure 2:**
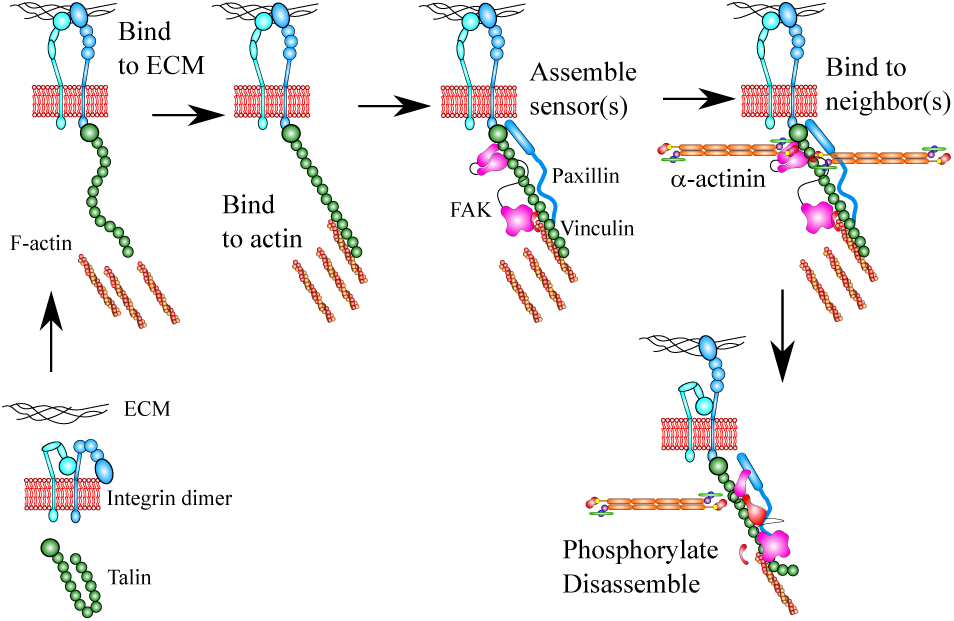
Sketch of a possible mechanism for the 5-step assembly of a mechanosensing protein complex Bell et al. (39). *α*-actinin is suggested as a possible basis for nearest-neighbour interactions between complexes. This implies that the sensor distribution can only start to aggregate after a large proportion of the sensors have formed, and are located close enough for the bridging protein to reach across to the neighbours.

If this explanation is accurate, the individual mechanosensors would have to be assembled some time before the cell reaches the hemispheric configuration at which it is observed to start binding to its neighbours (25). The discrepancy between the formation time of the individual sensor and of bound aggregates of multiple sensors is particularly important: if the sensor concentration was unstable at the time of its initial formation, small aggregates of integrin (nascent FAs) would already start to form before the cell reaches a benchmark hemispheric configuration. A mechanism for their aggregation would therefore require a geometry-related transition between a uniform and heterogeneous distribution. Fouchard et al. (25) suggest that the switch corresponds to the hemispheric configuration, but there does not appear to be any a priori reason for this observed criterion to be anything other than coincidental for the experimental situation which they considered.

If it is a real effect, this apparent time lag between the formation of the mechanosensor complexes and their aggregation into nascent FAs might fundamentally constrain the possible mechanisms for receptor aggregation in spreading cells. In particular, it has been suggested that this aggregation might occur via a kinetic mechanism, in which integrin-ECM binding bends the glycocalyx around an adhesion site and thereby creates a long range potential well within which mechanosensors can aggregate. However, if the receptor distribution were to remain stable for some time after the formation of mechanosensors, and if a kinetic aggregation mechanism were indeed to underlie receptor aggregation, either an increase in receptor concentration or a strengthening of the long-range interactions between receptors would be required to make the initially homogeneous distribution unstable. Both of these parameters could likely only increase smoothly and gradually, which would initially stop higher frequency modes from becoming unstable and likely lead to aggregation at low spatial frequency, contrary to the large number (order 100) of nascent FAs (25, 26).

As we shall see, this time lag between the initial formation of the individual mechanosensing complexes and their aggregation into nascent FAs naturally appears within our thermodynamic mechanism, which relies purely on short-range nearest-neighbour interactions. Should a short-range interaction be in play, one of constituents of the talin-bound cytoplasmic plaque would then likely allow for such an attractive interaction. *α*-actinin is a possible candidate for this as it is structurally symmetric and binds as a dimer to components of the protein plaque (54). The 36*nm* long *α*-actinin molecule (55) is definitely of the right order of magnitude length to mediate a direct interaction between mechanosensor complexes at short distances. Regardless of the nature of the molecular link (whether via *α*-actinin, or another multi-functional protein such as vinculin), such a direct short-range attractive interaction is only likely to occur at distances comparable to the length of an individual protein. This type of interactions can be modelled as simple nearest-neighbour interactions in a lattice gas model developed below.

### 1.3 Localisation of focal adhesion and geometrical simplifications

But where does the integrin-mediated cluster aggregation take place? Again, immunofluorescence observations of paxillin during cell spreading prove incredibly valuable (25, 26). It is found that paxillin – a component of the talin-mediated cytoplasmic plaque bound to integrin in the developed mechanosensor – is present at a high concentration very near the edge of the cell-ECM contact area after 3 minutes of spreading, that is, when the cell has spread to a near hemi-spherical shape. This in turn would suggest that integrins preferentially arrange themselves along the edge of this contact area. The true mechanism for this recruitment of integrin to the leading edge of the cell-ECM contact area is probably quite complex, but likely benefits from several cooperating mechanisms which help to determine both the spatial localisation of the adhesions and their temporal recruitment at the start of active spreading.

Classical arguments used in soft condensed matter can be used to show that receptor localisation in regions of high membrane curvature is energetically favourable as the energetic cost of bending the membrane is reduced by the presence of the receptor. The most highly curved region of the cell is clearly the edge of the cell-ECM contact area: on a 2D matrix, the rest of the contact area is flat, whereas the remainder of the cell membrane out of the ECM plane has an even larger radius of curvature than what was observed in the cell in suspension. This means in turn that by a succession of unbinding, fast diffusion and preferential binding to the edge of the cell-ECM contact area, we would expect an increase in integrin concentration at the edge of this area on a ring which we will henceforth dub the ‘*contact ring*’.

This likely takes place together with a mechanosensitive strengthening of individual mechanosensors which bind to the existing acto-myosin cortex located above the ECM plane. These are placed under increased load due to the contractility of the cytoskeleton, strengthening the integrin-matrix catch bonds and consequently increasing the residence time of the mechanosensor complexes within the thin intersectional area between the ECM plane and the out-of-plane cytoskeleton.

These mechanisms together may help to provide an explanation for the observed radial peak of the paxillin distribution near or on the contact ring as seen near the start of active spreading, which suggests that it might be possible to explore a highly simplified model of the distribution of mechanosensor complexes. To make initial progress towards such a model, we initially disregard the effect of the large-scale transient protrusive irregularities in the shape of the cell membrane which arise from the dissolution of membrane blebs and assume that the cell takes a shape such that it is under the same tension over its entire membrane. If the individual mechanosensing complexes have all formed before the cell has sufficiently spread, and if they preferentially form within the contact ring, then it is plausible to assume that the sensor concentration is very high along this ring at first.

In our attempt at the problem, we will assume that the concentration of mechanosensors is initially limited by the concentration of the most sparse but key cytoplasmic components of the mechanosensing complex, notably talin (56). Note that the synthesis of new proteins happens slowly relative to the spreading process (in tens of minutes to an hour), for the number of mechanosensor complex constituents to appreciably change throughout the initial phases of cell adhesion. Consequently, we will henceforth assume that the number of mechanosensors found within this contact ring is a constant number *N* once the cell has spread beyond a critical radius.

### 1.4 A note on turnover

Considerations of the aggregation of mechanosensors are complicated by the turnover of integrin-talin-FAK complexes (57–59): integrin opening under force allosterically changes the conformation of talin and subsequently also changes the conformation of FAK. FAK dephosphorylation irreversibly changes the conformation of FAK, breaking either the integrin-talin and the talin-FAK bonds (60). This detaches the mechanosensing complex from the integrin, but it is unreasonable to expect the entire complex to disassemble after this. More likely, it stays mostly intact and can bind anew to free inactive integrins. The probability of the mechanosensor’s cytoplasmic plaque being bound to integrin may then be determined in terms of the integrin-matrix binding rate, the integrin-talin binding rate, the stiffness-dependent integrin activation time and the FAK signalling time.

Regardless of the exact probability of the cytoplasmic plaque being bound to integrin, making the assumption that key parts of the mechanosensing complex remain intact even when talin and integrin are reverting back to their original conformations during the turnover process substantially simplifies the process. Indeed, for short-range interactions, if the molecule which is ultimately responsible for the nearest-neighbour interaction is bound to some of the proteins found in the region of talin, which is available to bind in both its closed and open configurations, then we would expect the binding strength between neighbouring mechanosensor cytoplasmic plaques to be roughly independent of whether or not they happen to be bound to the ECM via integrin. On the other hand, the kinetics of long-range kinetic aggregation mechanism would be seriously affected as the membrane deformation due to the presence of the mechanosensor would only come into effect when the sensor is fully assembled.

## 2 The model and methods

If two mechanosensing complexes are located sufficiently close that a protein can bind to each of them (within 30-50*nm* or so), then it is possible for a short-range interaction to be set up. If the concentration of sensors is sufficiently large, it would then seem likely that should such an interaction exist, it would dominate over the long-range potential-well set up by the distortion of the glycocalyx (47).

Any symmetric molecule found in the immediate vicinity of FAs in immunofluorescence imagery whose length is at least as long as the characteristic width of a mechanosensing complex is a prime candidate for this interaction. Given its availability close to the formation of the mechanosensors within the actin cortex as an actin cross-linker and due to its symmetric 36*nm* long dimer shape as well as its binding potential to vinculin and talin, *α*-actinin is a prime candidate for this interaction between integrin-mediated mechanosensors.

### 2.1 Thermodynamic mechanism for aggregation

The formation of nascent FAs (which takes place over the order of minutes) during cell spreading is a much faster process than mRNA transcription and protein translation (which take up to an hour in eukaryotic cells), but is much faster than diffusion across 1-10*µm* distances (given typical values for diffusion coefficient). It is therefore tempting to model the mechanosensor complex distribution within a cell as a thermalised distribution of a fixed number *N* of individual sensors, all located along the contact ring. The typical lifetime of the bonds to the sensor (20*s* or so for integrin-ECM bonds) complicates this picture a little, but should not be noticeable for aggregation times of the order of a few minutes.

Provided that the concentration of mechanosensors is highly peaked in the vicinity of the contact ring, one may assume that beyond a certain size of the contact radius, the vast majority of mechanosensor complexes are located upon the ring, and that their quantity within this thin area is in effect constant. We choose to model the contact ring between the cell and the surface where the response to extracellular forces is localised as a 1D lattice with cyclic boundary conditions. To replicate gradual cell spreading, we can adiabatically increase the length of the ring (or equivalently the number of equally-spaced lattice sites *L*). The situation is schematically drawn for the case of a one-mechanosensor thick ring in Fig. 3.

**Figure 3:**
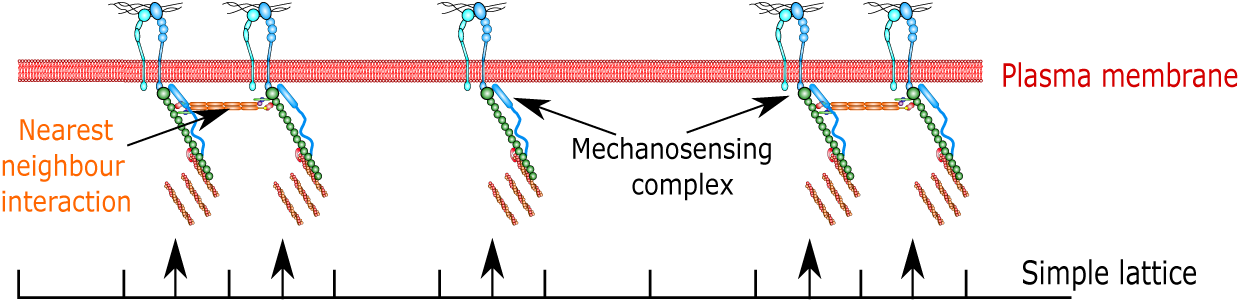
Schematic representation of the 1D lattice model of nearest-neighbour sensor interactions used to examine the aggregation of sensors along the contact ring between the spreading cell and the adhesive surface to which it is attached.

This sort of modelling is typical in a numerical approach, *e.g* Lepzelter et al. (61). An adaptation of the Ising model as considered by Guo and Levine (32) is the simplest possible model of the interaction (see sketch in Fig. 3):

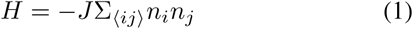

where Σ _⟨⟩_ indicates a sum over nearest neighbour pairs. The state of the sensor is indicated by the variable *n*, which in our case takes the values:

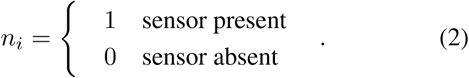

The binding energy of a single nearest-neighbour interaction is a constant *J* in this simple lattice model, whose exact value likely depends on cell type. With these simplifications in hand, we will find the Ginzburg-Landau action (which is proportional to the free energy of the focal adhesion complex distribution) and examine the spatio-temporal evolution of a the distribution of force sensing complexes located on the contact ring between the cell and an adhesive surface.

### 2.2 Sensor interaction model

For any process which occurs on a much shorter timescale than cell spreading (seconds rather than minutes), the length of the contact ring between a cell and an adhesive surface can be assumed to be constant. Modelling this ring as a lattice gas with constant site spacing, the number of sites *L* is constant over such a short time interval. Using this simplification, we can closely mirror the derivation for the classical Ising model over a sufficiently short time interval.

The single-molecule partition function for the lattice gas model with Hamiltonian *H* from Eqn.(1) is:

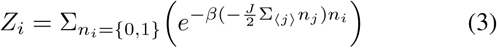

where the sum runs over all of the *L* sites around the ring, and *β* = 1*/k*_*B*_*T* as usual.

The full partition function is the product of *L* partition functions for each site, subject to the constraint *N* = Σ_*i*_*n*_*i*_ of the constant total number of individual sensor complexes. This condition can be expressed with a delta function, and rearranged into matrix form as:

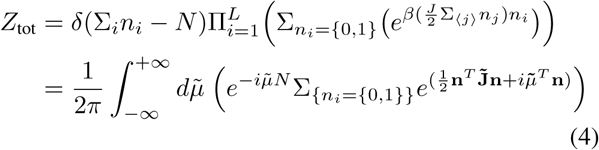

where we have defined the modified non-dimensional coupling constant:

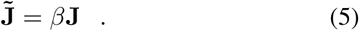

Using the *N* -dimensional Gaussian integral identity, we can write the second exponential term in Eqn.(4) as:

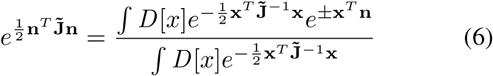

where **x** is the Hubbard-Stratonovich vector field, which we need to later obtain the order parameter description. Note the ambiguity in the sign of the term whose exponent is linear in **x** in Eqn. (6); both formulations will later be shown to be identical when expressed with respect to the physical focal adhesion complex density.

Treating **x** as independent of **n**, we can factorise the partition function (4) as follows:

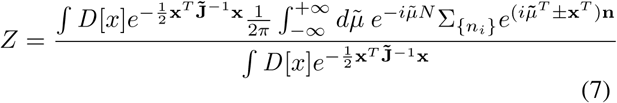

The integral of the linear part of Eqn.(7) can be simplified:

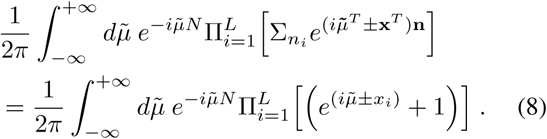

An action functional 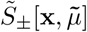 can now be defined following the steps of Landau-Ginzburg-Wilson theory:

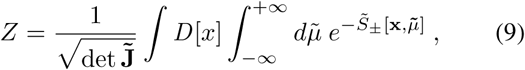

and found to be in the most general form:

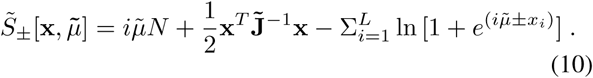

The next step is to calculate the expectation value of the Hubbard-Stratonovich field:

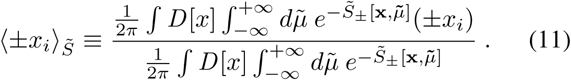

After introducing an auxiliary *N* -component column vector **y** = (*y*_1_, …, *y*_*N*_)^*T*^, we can re-express the Hubbard-Stratonovich field components as:

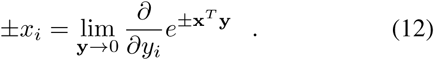

Then performing Gaussian integration in Eqn.(11) modified with *n* → *n* + *y*, we obtain 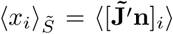 if the sign of the exponent of the linear term 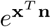 in Eqn.(6) is chosen to be positive; or 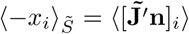 if this exponent is instead negative.

We need a variable whose expectation value can be identified with the average activation per site, and so define the order parameter:

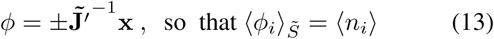

This identifies the discrete field *ϕ*_*i*_ as the local mechanosensor concentration at site *i*. We will later transform this to a continuous density *ϕ* which depends on a continuous position along the adhesion contact ring.

An immediate consequence of the definition of the physically meaningful density field *ϕ* in Eqn.(13) is that the previously noted sign ambiguity is lifted: both formulations lead to identical expressions when expressed in terms of *ϕ*. We may therefore without loss of generality only consider the case where the exponent of the linear term 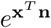 in (6) is positive.

It is convenient at this stage to define the fluctuation of the Hubbard-Stratonovich field over all *L* lattice sites: 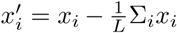. The corresponding fluctuation of the concentration field about its average value *ϕ*_*a*_ = *N/L* is directly related: 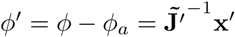.

At equilibrium, the action is maximum with respect to 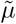, or equivalently with respect to the modified variable 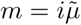 (according to the condition of stationary phase). Carrying out this differentiation of Eqn. (10) allows us to express *m* in terms of the Hubbard-Stratonovich field **x** and the system variables *L* and *N* in equilibrium:

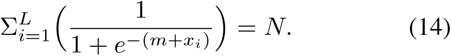

The Ginzburg-Landau action and its variable constraints have thus far been expressed in an exact, but rather unwieldy form. We will have to make several strong approximations in the next section to proceed towards a manageable form for *S*.

### Key approximation 1: comparison of the occupancy of full and empty states

Near the start of spreading, we assume that the majority of sites are full on the contact ring of an initially small footprint, so *N* can be assumed to be the same order as *L*. This means that the probability of a single site being in the ‘filled’ state (i.e. contain a mechanosensor complex) is much greater than that of being in the ‘empty’ state. This gives the following condition for the Boltzmann factors:

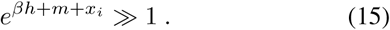

When (*L*−*N*) ≪ *L*, we can simplify the constraint (14) and express it in a form which makes its solution possible:

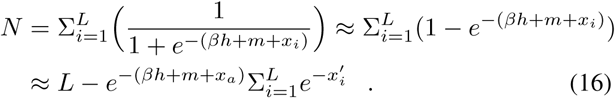

This can be rearranged to:

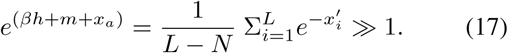

As expected, this is a large quantity if the number of sites and the number of sensors are both large.

### Key approximation 2: series expansion for small nonuniformity

Using this approximation, we perform the series expansion of the action in terms of the Hubbard-Stratonovich field **x**, details in Appendix A. It is only valid when **x** is small, which we will have to re-examine later. In Appendix B, the partition function and the effective action *S*[*ϕ*] are rewritten, transformed into Fourier space, made continuous, and finally transformed back into real space. We find a form of action, which in this context could be called the effective free energy, that has recognizable features of the Ginzburg-Landau theory, where we retain cubic and quartic terms in the order parameter expansion:

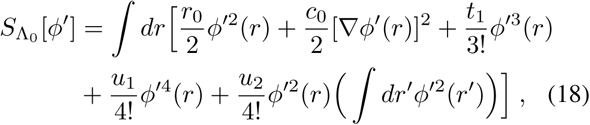

where *ϕ*′(*r*) is the continuous density of fluctuations about the cell-averaged density *ϕ*_*a*_ = *N/L*, and the coefficients are explicitly defined in Appendix B. Specifically, the two quadratic-order coefficients take the form

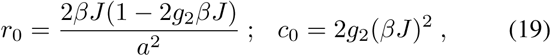

with

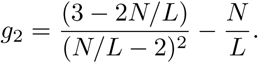

We see that the control parameter (replacing the temperature in the classical theory of phase transitions) is the ratio *N/L*, which starts at 1 when the cell adhesion ring first forms, and then diminishes as the cell spreads and the ring perimeter *L* grows. The gradient coefficient *c*_0_ remains positive, but the ‘main’ coefficient *r*_0_ could become negative (thus causing the instability of the homogeneous density distribution around the adhesion ring) at a critical value of *N/L*, which depends on the coupling strength *βJ*. At large *βJ* the instability occurs almost immediately below *N/L* = 1, while for *βJ* < 1 the coefficient *r*_0_ remains positive for all *N/L* and no instability occurs, see Fig. 4.

**Figure 4:**
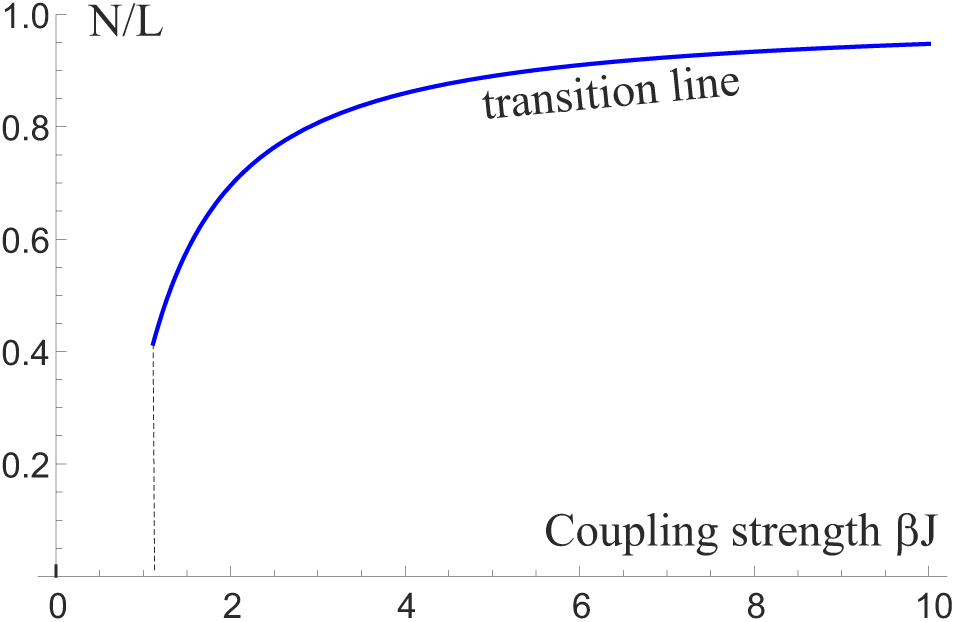
The ‘phase diagram’ of values of *N/L* vs. the sensor interaction strength *βJ*, showing the transition line at which the coefficient *r*_0_ from Eqn.(19) changes sign, and the azimuthal instability becomes possible. *r*_0_ remains positive for *βJ* smaller than the critical value close to 1, as labelled on the plot.

In Appendix A, we see that the coefficients of terms in the series expansion to the *n*^th^ power in the Hubbard-Stratonovich field **x** are of order unity, allowing for accurate results while fluctuations about the average field are small.

## 3 Results and comparison with experimental data

### 3.1 Spatial frequency of fluctuations

In the immediate vicinity of the critical point, the quadratic-order terms in the density of sensor concentration fluctuations *ϕ*′ are much larger than all of the higher order terms. The action (effective free energy) written in Eqn.(18) can therefore be substantially simplified in the vicinity of the critical point:

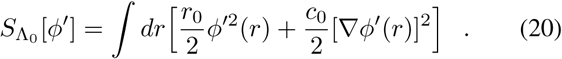

The time-dependence of the concentration fluctuation can be described by the Cahn-Hilliard equation (62), which near the critical point takes the form:

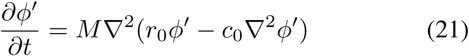

where *M* is a (constant) mobility, to be experimentally determined. Its behaviour is subject to the boundary conditions:

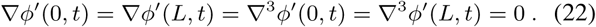

A well-known solution satisfies these boundary conditions:

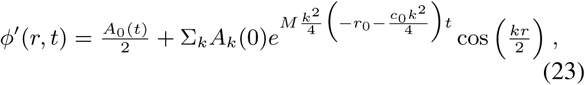

where *k* is the wavevector, directly related to the number *m* of peaks in the azimuthal modulation of sensor concentration: *k* = 2*πm/L*.

The fastest growing wavenumber, which corresponds to the oscillation lengthscale that maximises the exponential term above, is:

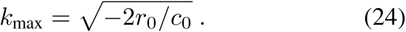

The critical mode number, for the largest spatial frequency at which the exponential term could grow with time is

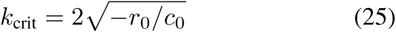

(see Eqn.(19) and Appendix B for the values of *r*_0_ and *c*_0_, and Fig. 4 showing when *r*_0_ becomes negative).

If we now also include periodic boundary conditions: *ϕ*′(*r, t*) = *ϕ*′(*L, t*) and ∇^2^*ϕ*′(*r, t*) = ∇^2^*ϕ*′(*L, t*), terms with odd *m* in the solution for *ϕ*′(*r, t*) are eliminated. To see a single mode grow, we therefore require that

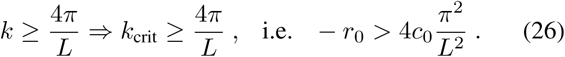

Given a fixed number of force sensing complexes *N* on the contact ring, and a specific interaction energy between neighbouring complexes, this condition allows us to determine the minimum value of *L* at which the cell stops being azimuthally uniform and starts to form spatial patterns around the contact ring, as illustrated in Fig. 5. But this in itself would not be particularly useful: lower modes take a very long time to grow due to diffusive constraints. The important question is rather whether we can identify a mode which grows sufficiently fast that it dominates before subsequent cell spreading allows for higher frequency modes to become unstable and then dominate. This exploration will be made easier by comparing with rough physiological values.

**Figure 5:**
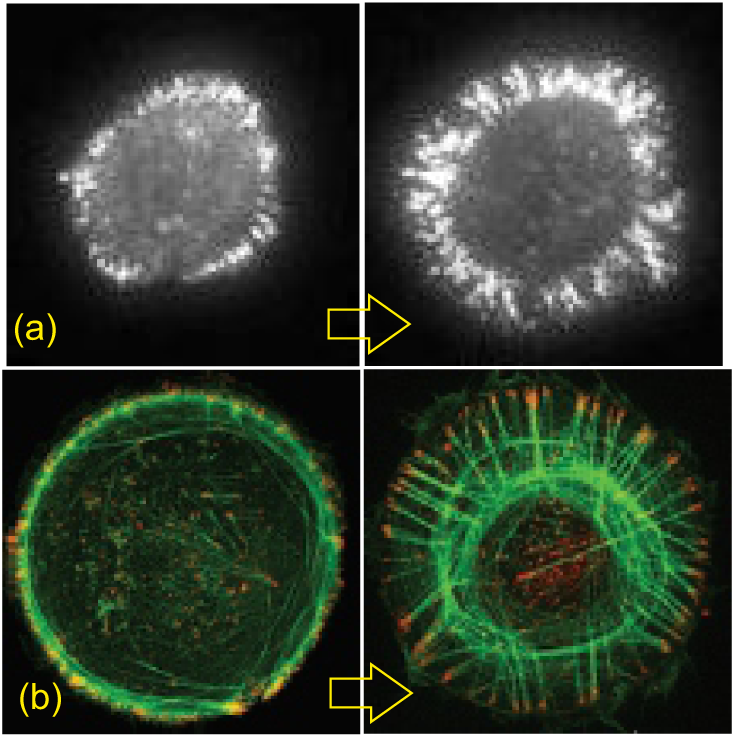
The transformation of density of adhesion complexes (labelled by fluorescence of paxillin) as the contact ring expands on cell spreading. Images (a) are from Fouchard et al. (25), and images (b) are from Tee et al. (26), with permission.

### 3.2 The effect of spreading on nascent FA number and comparison with experiments

As mentioned above, observations by Étienne and Duperray (40) indicate that the cell-ECM contact area increases noisily during cell spreading; consequently, the contact ring must be expected to increase via a process of significant jumps and periods of stagnation or retraction, so *L* must increase noisily and abrubtly. A possible driving force for this is the aforementioned disassembly of membrane blebs required for the maintenance of tension as well as membrane and cytoskeletal material throughout the spreading process.

Observations by Norman et al. (63) show that these blebs take on a variety of shapes and sizes, but a rough estimate suggests that each individual bleb has a radius which is of the order of one twentieth to one tenth of the radius of the cell as it comes in contact with the surface. Both structures are approximately spherical, so the area of one bleb would be approximately 1*/*400 < *A*_bleb_*/A*_cell_ < 1*/*100 of the otal area of the cell. When the cell is in the hemispherical configuration, releasing a bleb of this kind – assuming that the actin that it contains is spread out equally throughout the cytoskeleton – would result in an increase in cell radius of the order of 1*/*800 < Δ*r/r*_cell_ < 1*/*200 of the cell, i.e. increase the lattice size *L* by this proportion. Now, nascent FA formation takes place initially when the radius of the cell is *r*_cell_ ≈ 15 − 20*µm* for fibroblasts. So assuming a lattice spacing of *a* ≈ 40*nm*, we obtain an estimate of *L* = 2*πr*_cell_*/a* ≈ 2000 − 3000. The increase in lattice size due to the disassembly of the bleb would then be Δ*L* = (Δ*r/r*_cell_)*L* ≈ 3 − 15.

But according to the above reasoning, for a constant number of sensors *N*, the larger the lattice size *L*, the higher the fastest growing mode is. Provided that the spatial modes grow sufficiently fast, we therefore expect to find that the typical size of mechanosensor aggregates is related to the root mean square (RMS) value obtainable from the membrane bleb size distribution if this mechanism is indeed correct. This is one step beyond our current work, and so for simplicity we will assume that the membrane bleb size distribution is sharply peaked about its average value. Regardless, it is important to note that we expect there to be some substantial stochasticity to the actual number of nascent FAs in spreading cells, and we suggest that a possible means of distinguishing between a kinetic and a thermodynamic aggregation process might involve exploring the shape of the observed FA frequency distribution in a population of cells.

In arriving at an order-of magnitude estimate for the spatial frequency of nascent FAs, illustrated in Fig. 5, we consider the limiting case in which the mechanosensor concentration is at its critical point prior to the bleb merging with the cell: in mathematical terms, the coefficient *r*_0_ = 0 and no spatial modes are unstable. This then gives a (complicated) equation for *L*_critical_ in terms of *N* and the coupling energy *J*, which is plotted in Fig. 4 above. As *L*_critical_ + Δ*L* is then substituted back into the equation for the fastest-growing mode *k*_max_, we find estimates for the upper bound for *k*_max_ for typical physiological values corresponding to the situation observed by Fouchard et al. (25), which are presented in a ‘phase diagram’ Fig. 6

**Figure 6:**
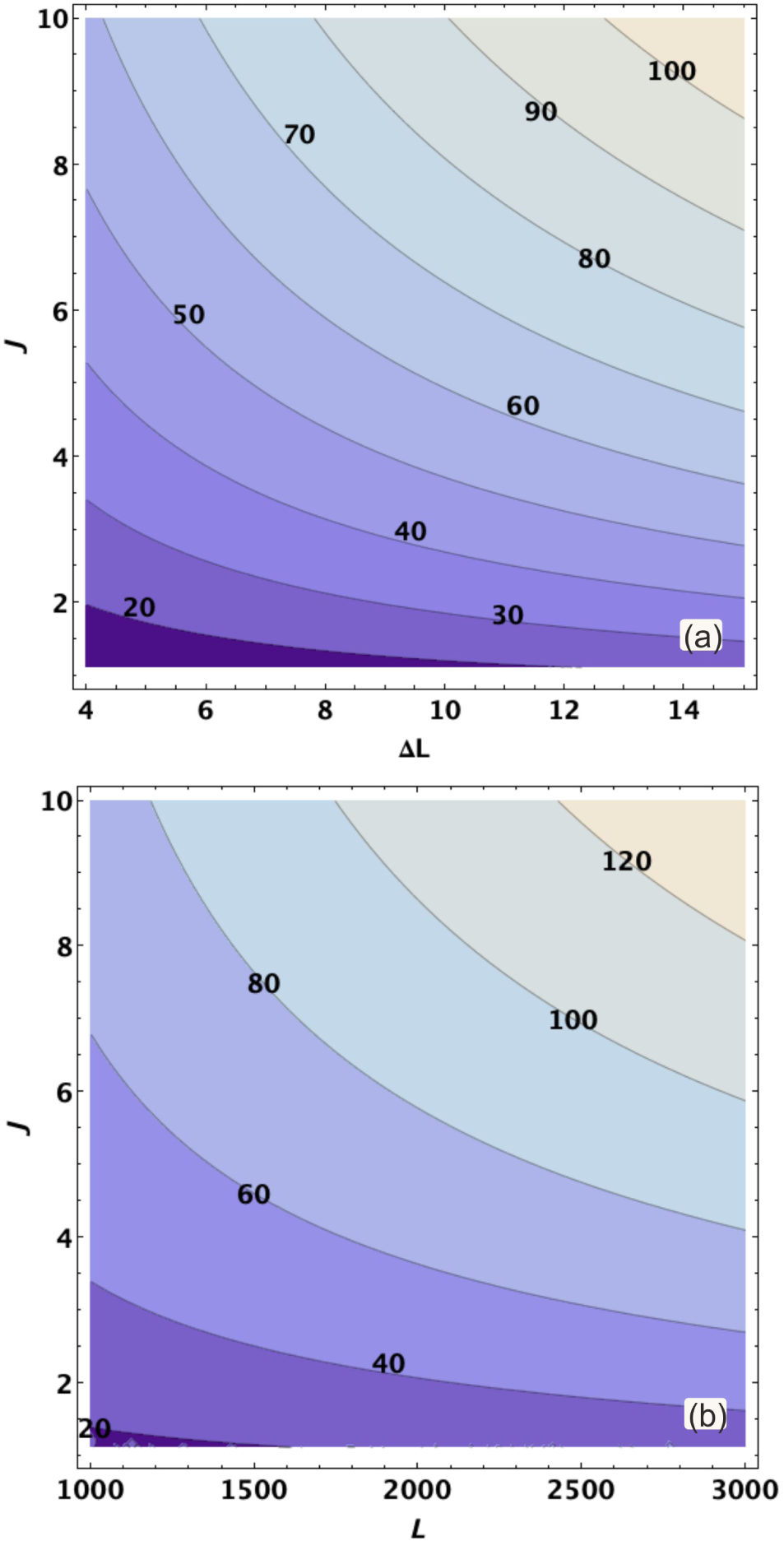
Contour plots of the fastest growing mode *m*_*max*_ = *k*_*max*_*L/*2*π* found in our lattice model if the cell spreads by an amount Δ*L* from a critical length *L* by rapid bleb retraction. (a) At *L* = 2000 corresponding to a spread radius of *r*_spread_ ≈ 15*nm* if the typical size of a mechanosensor is *a* ≈ 40*nm*. The contour map is a function of the change in lattice length Δ*L* and the coupling strength *J*. (b) At Δ*L* = 15, which corresponds to the increase in lattice length when a large bleb retracts into the body of the cell. Contour map is a function of the critical lattice length *L* and coupling strength *J*, given in units of *k*_*B*_*T*. Assuming that the bond strength is similar to that of the *α*-actinin to actin bond, which has been reported at 2*k*_*B*_*T* by Miyata (64), the zone of interest for nearest-neighbour mechanosensor interactions would be in the bottom half of the graph.These results are in no way meant to be quantitative, but altogether strongly suggest that the spatial frequency of nascent FAs is in the range of *m* ≈ 20 − 100. This is of the same order of magnitude as the number of nascent Fas

### 3.3 Mode growth time constraints

The number of nascent FAs may also be strongly constrained by the mode growth time, which is found for classical spinodal decomposition to be proportional to the second power of the pattern wavelength *λ*_max_, or alternatively 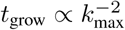. Indeed, prior to aggregation, mode growth time should be expected to be proportional to the single-cluster diffusion time. The prefactor is of order unity and can be found by substituting a regular solution with a particular scale width into the free energy. A standard reasoning gives:

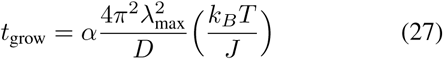

where *α* is a constant of order unity.

Integrin diffusion coefficients have been found to be as high as *D*_integrin_ = 10^−1^*µm*^2^*s*^−1^ for free integrin (65). Assuming that the mechanosensing cluster diffusion time is comparable to the integrin diffusion time, we may numerically evaluate the above expression with *λ*_max_ = 2*πr*_spread_*/m*_max_ for a typical *r*_spread_ ≈ 15*µm* at the time of the formation of nascent adhesions. Bearing in mind that in doing this we have likely overestimated the mechanosensor diffusion coefficient by a substantial amount, we can use this to obtain a lower bound for the nascent FA aggregation time.

In particular, we find that low mode numbers (order 10) take much longer than the full spreading time (≈ 30mins) to grow in this way, and so cannot lead to the formation of the observed nascent FAs. Thus, the cell must spread sufficiently beyond the critical point *r*_0_ = 0 to allow for higher frequency modes to become unstable. This can be achieved either if a large bleb is disassembled or if several blebs are successively assimilated into the cell body.

Cell spreading takes place relatively rapidly with a ball-park radial expansion speed for the cell-ECM contact area of ≈ 1 *µm/min* near the start of active spreading. Following the above estimates for bleb size, this corresponds to an average expansion rate of the lattice of Δ*L* ≈ 25*t*, with the time *t* expressed in minutes. A point of comparison can be made between the fastest growing wavelength at time *t* and the mode whose growth time corresponds to the disassembly time of a particular bleb. The result of this is that the nascent FA spacing *m*_max_ would be of order 100 for the nascent FA to aggregate before the cell substantially spreads further. One might argue that this is seen in experiments illustrated in Fig. 5.

We have therefore identified both a lower bound and an upper bound to the number of nascent FA which develop at the start of spreading, and this appears to be generally in the vicinity of 100. This number can change over time however: once the unstable modes have grown out, the pattern coarsens according to the classical theory of spinodal decomposition (62). At the later stages of spinodal decomposition, and the beginnings of coarsening, the initial aggregation length scale becomes less clear (66). In the classic Cahn-Hilliard coarsening, the phase boundaries are diffuse as the initial length scale still dictates the width of the domain walls. FAs on the other hand do not appear to be diffuse at all, but this is not necessarily an issue as the cell keeps on spreading, thereby increasing the frequency of the fastest growing mode even further and narrowing the FA boundaries.

## 4 Discussion and conclusions

An analytical physical approach has already proven highly useful and insightful in exploring the problem of cell spreading by accounting for the vertical and radial dynamics of the cell shape throughout the process. Here we examine the question of the development of azimuthal structure and evolution of a spreading cell when viewed from above from a circular shape to an anisotropic one.

Recent biological data lends credence to the idea that the aggregation of integrin-mediated mechanosensors found in focal adhesions is responsible for the development of initial azimuthal inhomogeneity. In particular, we suggest the possibility that a thermodynamic mechanism relying on short-range protein bonding plays a substantial role in cell spreading, and identify the ring at the intersection of the cell basal plane and the out-of-plane acto-myosin cortex as a possible location for the recruitment and the aggregation of its constituent proteins.

This thermodynamic mechanism may potentially cooperate with a kinetic mechanism of receptor clustering which would rely on long-range mechanosensor-induced membrane deformations to attract other bound integrins. Although both mechanisms involve very different interactions, it is not immediately clear how to distinguish their predictions – not only because of the lack of accurate biological data, but because many of these are similar: both predict that mechanosensors will aggregate on a particular length scale for instance.

One possibility for testing their differences would be to more accurately measure the onset time of the initial aggregation of mechanosensors. If this time does not happen to coincide with a particular cell shape (*e.g.* a quasi-hemispherical cell), then this would lend credence to the idea that a switch other than a purely geometrical one is required, favouring the thermodynamic model. To test the validity of a kinetic mechanism in a particular situation, it might be possible to measure the membrane deformation stresses around bound integrins in order to determine the strength and width of the kinetic well.

Regardless of the specific mechanism in play, the aggregation of integrin-mediated mechanosensors into clusters has clear consequences for the subsequent evolution of the cell’s shape during the remainder of spreading. The formation of focal adhesions coincides with the extension of the lamella via F-actin polymerisation. Newly formed F-actin filaments push against ECM-bound nascent FAs and are able to exert a protrusive force on the membrane, which in turn leads to further cell spreading and sets up a retrograde flow of actin within the cell behind the adhesions. This in turn likely helps set up a clear radial directionality to the growth of FAs, which become long and radially-oriented. Therefore, while a 1D model was useful in examining the initial aggregation of mechanosensors at the start of the active spreading process, it is insufficient to understand their growth and decay towards the later stages of cell spreading. Nonetheless, observations by Fouchard et al. (25) clearly show that the angular distribution of the initial pattern which we investigated in this work correlates strongly with the subsequent distribution of FAs. An obvious next step towards understanding the clustering of FAs during differentiation and movement would be to more accurately model FA maturation, decay and interactions between neighbouring clusters.

Finally, we suggest that a similar contact ring might arise on either side of a cell-cell junction between two interacting eukaryotic cells. A similar mechanism might lead to cadherin aggregation into adherens junctions after cadherin-actin bonds have developed (67, 68). This might have an even more biologically relevant role, as the adhesion surface between two cells is much more naturally two-dimensional than an in-vivo ECM surface.

## Author contribution

Both authors conceived the idea, carried out different elements of data analysis and wrote the paper.

## Acknowledgements

The authors acknowledge many helpful discussions with Sam Bell, Theresa Jakuzeit, and Heidi Welch, as well as J. Fouchard, A. Asnacios, and A. Bershadsky for giving access to their experimental results. This work has been funded by BBRSC DTP Cambridge (grant no. EP/M508007/1).

## Appendix A: Series expansion of the action

By explicitly substituting 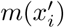 as obtained in Eqn. (14) into Eqn. (10), we can obtain a useful expression for the action 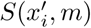 in terms of 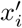 only. Doing so mathematically incorporates the *N* = const. constraint on the number of mechanosensing complexes into Ginzburg-Landau action and will allow us to subsequently formulate it in terms of the physical sensor concentration fluctuation *ϕ*′. We begin by writing the exponential 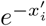 as a series:

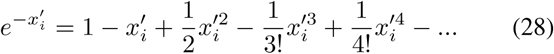

Summing over all of the states *i* gives:

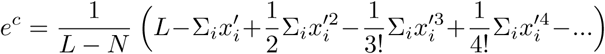

To keep track of the order of fields, we introduce the parameter *a* through:

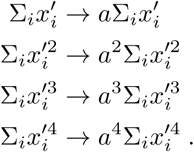

We now expand the above in powers of *a*:

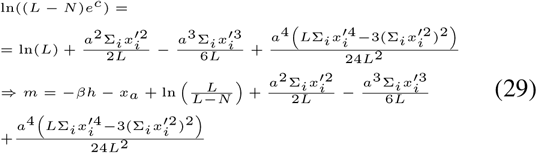

A similar trick can be used to keep track of the order of the terms in the expansion of 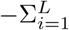 In 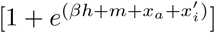, substituting in the expression for the Boltzmann factor. We are interested in the expansion of:

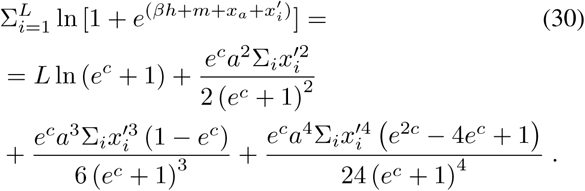

Substituting in the expansion of the Boltzmann factor *e*^*c*^, we find the series expansion of this term up to 6^th^ order in *a*:

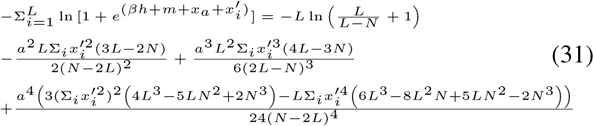

Setting the series parameter *a* = 1, we forget for the time being about the explicit dependency of the terms on *L* and *N*, and write:

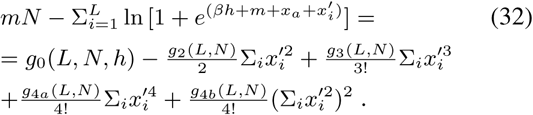

where we make the identifications:

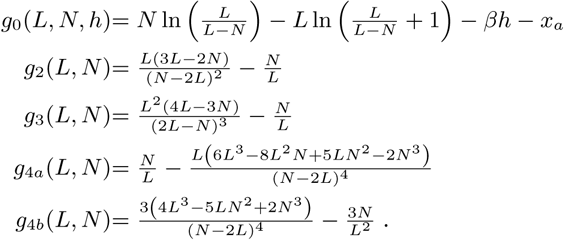

The action (or the effective free energy) can be reexpressed in terms of the average value of the Hubbard-Stratonovich field in order to substitute the above:

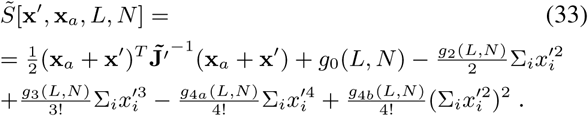

We will also need to check that the total contribution of the sixth order term is positive in order for the Hamiltonian to be bounded from below.

## Appendix B: Reexpressing the action in terms of a continuous sensor concentration

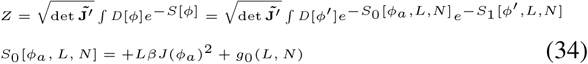

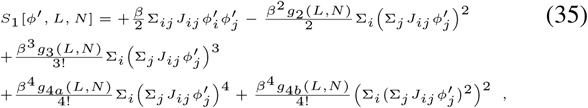

where the couplings *J*_*ij*_ of the original theory have been restored.

The order parameter can be transformed into wave vector space, imposing periodic boundary conditions:

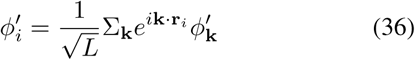

where the wave vectors are quantised as 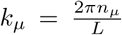 with *n*_*µ*_ = 0, 1, …, *N*_*µ*_ − 1 where *N*_*µ*_ is the number of lattice sites in the direction *µ* (this naturally allows the problem to be extended…) s.t. 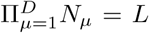 is the total number of lattice sites. We use the identity:

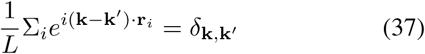

to obtain the Fourier transform of the terms in the truncated effective action:

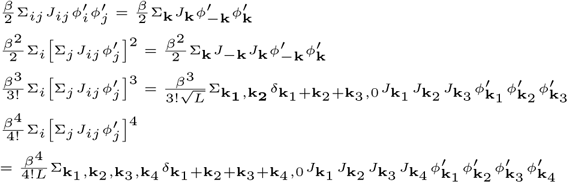

where *J*_**k**_ is the Fourier transform of the exchange couplings *J*_*ij*_ ≡ *J*_(_**r**_*i*_ − **r**_*j*_):

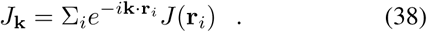

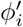 and *J*_*ij*_ are real, and the couplings are reciprocal: *J* (−**r**) = *J* (**r**), so 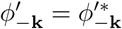 and *J*_−**k**_ = *J*_**k**_.

Thus in Fourier space, the truncated effective action may be written:

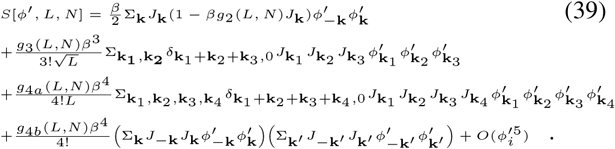

Sufficiently close to the critical point, only long-range fluctuations (so **k** small) contribute significantly. So *J*_**k**_ can be expanded in powers of **k**. If the coordination number is *z* = 2*D*, then:

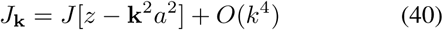

So the quadratic term in the field *ϕ*′ has the following coefficient:

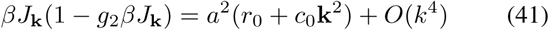

where |*T* − *T*_*c*_| ≪ *T*_*c*_ is assumed (so the coefficient of the quadratic term in *k* simplifies) and the important constants *r*_0_ and *c*_0_ are defined as:

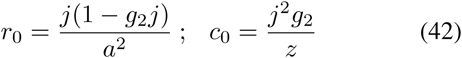

where *j* = *zβJ*.

In the limit of infinite volume *V* = *La*^*D*^ → ∞, the discrete set of allowed wave vectors merges into a continuum: replace the momentum sums by integrations according to:

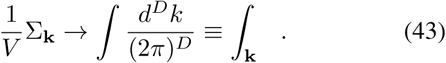

Normalising the fields by the continuum fields 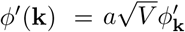 sinstead, we define the coupling constants:

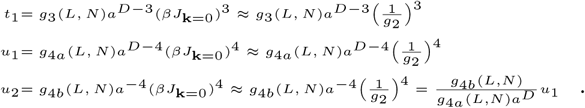

In the main text, we treat the evolution of a sensor concentration on the adhesion contact ring between a cell and a surface as a 1*D* problem. It is clear that in 1*D* (with *D* = 1 by definition), if we set *V* = *L*, then *a* = 1 and the above constants simplify considerably. Wavevectors may also be rewritten without any confusion in scalar form.

To check the dimensionality, remember that in this situation, there are in effect not four but three integrals, as one is eliminated in the evaluation of the delta function. The effective action becomes (note, there is an ultraviolet cut-off to the integration):

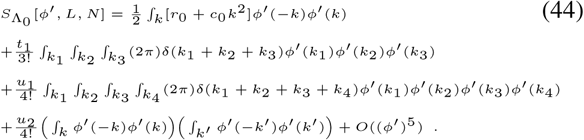

Transforming into real space, we obtain another form for the action:

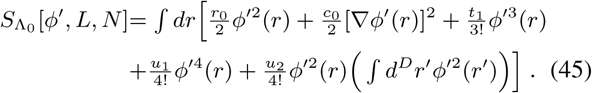

## Appendix C: Behaviour away from the critical point

The action in Eqn.(18) is hard to analyse without making some simplifying assumptions. We saw in the previous section that fluctuations in the sensor concentration are well-described by a set of sinusoidal perturbations with a given wavenumber *k* near the critical point. We can examine each of these contributions separately while fluctuations are still sinusoidal (*i.e.* provided that the cubic, quartic and higher order terms in the action do not dominate over the quadratic term), considering:

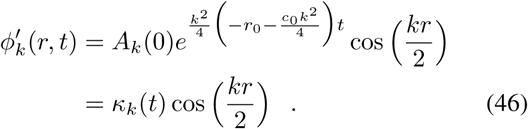

In the long-time limit, the mode closest to the critical mode will dominate, and so we will approximately find that:

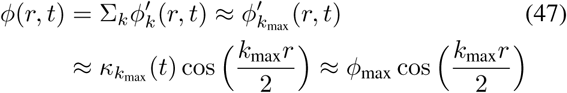

where *ϕ*_max_ is the amplitude of fluctuations. This enormously simplifies the problem, as the integral in the fourth order term can be approximated as:

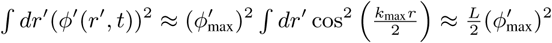

The Ginzburg-Landau action can then be written in this limit as:

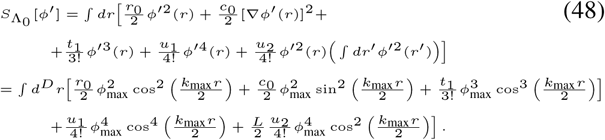

Now cos^2^ *θ* ≥ cos^4^ *θ*, ∀*θ*, so if we show that *u*_2_ > 0 and that the fourth order terms in the action above are positive for *k*_max_*r* = 2*mπ*, for integer *m*, then they will be positive at every point *r* along the boundary of the cell.

Using the results from Appendix B, we can show that when *k*_max_*r* = 2*mπ*, the fourth order terms combine to:

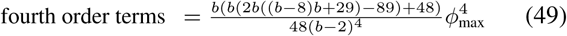

where *b* = *N/L*. This combination is positive ∀*N* < *L*, so we expect the fourth order terms to be positive at all positions *r* along the cell boundary, for all sensor concentrations for which our Boltzmann factor approximation is appropriate (so *L* − *N* ≪ *L*) and when the modes have grown for a sufficiently long time that the mode closest to the fastest growing spatial frequency *k*_max_ dominates.

In this limit, a fourth order expansion of the Ginzburg-Landau action is therefore sufficient to describe the destabilisation and initial growth of spatial modes (by analysing the quadratic terms) as well as the development of two slightly asymmetrical minima in the action profile (due to the presence of non-zero third order terms) relative to the cell-wide average sensor concentration. This behaviour is examined in Fig. 7.

**Figure 7:**
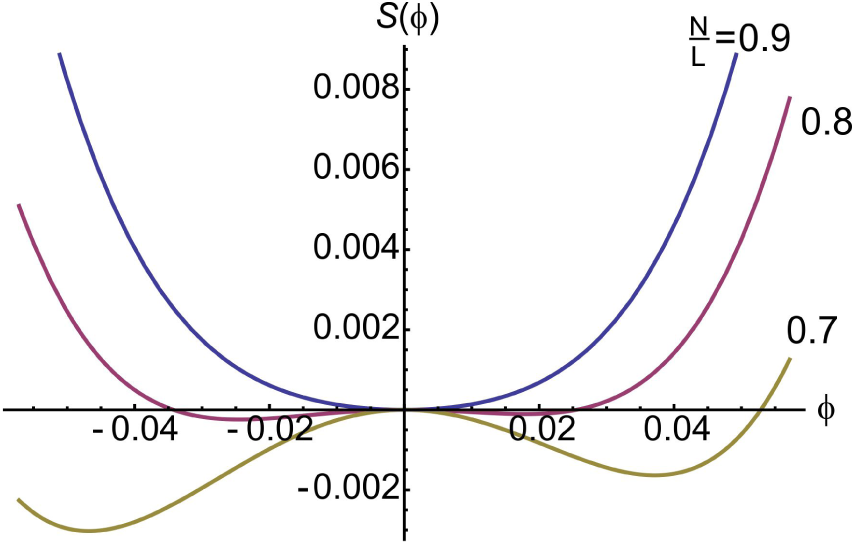
Plot of the profile of the Ginzburg-Landau action as a function of the amplitude of periodic deviations from the average sensor density as the average sensor density decreases. A nearest-neighbour binding constant of *J* = 3.5*k*_*B*_*T* is used as an illustrative value. The action is plotted at the spatial *r* positions along the cell-ECM contact rim which correspond to the peaks in the sensor distribution as these are found to produce the most asymmetric action landscapes. As the cell spreads, *N/L* decreases and the *ϕ* = 0 configuration (average concentration) transitions from a global minimum to a local maximum of the action, and two minima develop on either side of the average sensor concentration; the system undergoes a phase transition from a uniform sensor distribution to a two phase distribution in which a denser and a more dilute phase coexist. The profile of the action is asymmetric due to the presence of a non-zero third order term. This term is small and it is possible that the asymmetry in the action might be attributable to the error in the Boltzmann factor approximation introduced in Eqn. 15 as the condition that *N* − *L* ≪ *L* is no longer verified for smaller values of *N/L*.

